# Resilience and reassembly of redox-structured microbial functional networks after rare holomixis in a meromictic lake

**DOI:** 10.64898/2026.01.18.698864

**Authors:** Ya-fan Chan, Pei-Wen Chiang, Sim Lin Lim, Denis Rogozin, Vladimir Zykov, Chong-Heng Ye, Sen-Lin Tang

## Abstract

Land-locked meromictic lakes are characterized by long-term stratification and steep redox gradients that sustain vertically structured microbial communities and tightly coupled biogeochemical processes. Because complete overturns of the lake are rare, the dynamics of microbial reassembly after redox gradients collapse and subsequently recover remain poorly resolved. We investigated a mixing–restratification transition in Lake Shira (Siberia) with depth-stratified sampling of oxic, chemocline, anoxic, and water–sediment interface layers across four stages: intermittent holomictic (IH), complete holomictic (CH), developing meromictic (DM), and stable meromictic (M). 16S rRNA gene amplicons showed that CH homogenized the water column, dominated by *Gamma*- and *Alphaproteobacteria*, *Campylobacteria*, and *Cyanobacteriia*. As stratification re-formed (DM–M), communities became strongly depth-partitioned, with *Desulfobacterota* and other anaerobes re-established in sulfidic deep waters and assemblages concentrating near the redox transition. Nanopore metagenomics reconstructed 401 MAGs, revealing stage- and depth-specific functional repertoires consistent with redox zonation. Core MAGs, including *Yoonia* spp., persisted across phases, suggesting functional continuity underpinning rapid ecosystem recovery. These data provide a system-wide view of biogeochemical reassembly during collapse and restoration of stratification meromictic lake.

## Introduction

Meromictic lakes are distinguished by their permanent or long-time stratification, where the water column does not mix fully, leaving a high density, isolated bottom layer separate from the more dynamic surface waters. This stable physical stratification produces a persistent vertical redox gradient, from an oxygen-rich upper layer through a steep chemocline (transition zone) into an anoxic, typically sulfidic, deep layer (*1*). Microbial communities in these lakes are highly structured and constituted by depth, aligning with sharp gradients in light, oxygen, nutrients and reduced chemicals (*2, 3*). The upper mixolimnion supports oxygenic phototrophs and aerobic heterotrophs, whereas the anoxic monimolimnion hosts strictly anaerobic metabolisms such as sulfate reduction and methanogenesis (*4–6*). The chemocline, in particular, often concentrates dense populations of anoxygenic phototrophs that thrive opposing gradients of light and sulfide (*7*). For example, the chemocline of Lake Cadagno in Switzerland harbors a compact layer dominated by purple sulfur bacteria (PSB) and green sulfur bacteria (GSB) that oxidize sulfide anoxygenically (*5*). Similar enrichments occur in meromictic Fayetteville Green Lake in USA, the PSB, GSB, and anoxygenic photosynthesis-capable cyanobacteria were both abundant within the chemocline zone (*3*), and three meromictic lakes inn Asia (Lakes Shira, Shunet, and Oigon), where chemocline communities were mainly dominated by PSB (*4*). Thus, each stratified layer in meromictic lakes harbors a distinct microbial assemblage. And, the dominant taxa and functional groups within each layer can differ substantially among lakes, reflecting lake-specific geochemistry, light climate, and basin characteristics. For decades, some meromictic lakes like Cadagno Lake, Green Lake, Lake Shunet and Ace Lake, have become model ecosystems for studying microbial ecology under stable redox stratification (*3, 5, 8, 9*).

Although meromictic lakes are physically stable, they are not fully resistant to environmental disturbances. Climate variability and episodic events can perturb the stratification, impacting the lake’s microbial diversity and ecosystem processes (*1*). Previous studies revealed that the stability of meromictic lakes can be disrupted by various abiotic factors, such as wind-driven circulation, freshwater inflows, changes in water level, or extreme weather events. From example, The Lake Lugano North Basin has been meromictic for several decades, with anoxic waters below 100 m depth. Two consecutive cold winters in 2005 and 2006 induced exceptional deep mixing, leading to a temporary oxygenation of the whole water column (*10*). With the ventilation of deep waters and the oxidation of large quantities of reduced solutes, the lake’s total redox-balance turned positive. Shifts of climate and hydrology ever can alter the mixing depths of meromictic lake. Increased precipitation or freshwater runoff can be increased stratification; in contrast, prolonged warming can be decreased density gradient and stratification by reducing the physical or chemical differences between the mixolimnion and monimonlimnion (*11, 12*). Additional studies revealed that, when meromixis breaks down, isolated nutrients from the monimolimnion are introduced into the upper layers, stimulating ecological shifts across microbial, phytoplankton, and zooplankton communities (*13–15*). However, such complete overturns are extremely rare in meromictic systems as the strong density gradients usually prevent full circulation.

Lake Shira, a saline lake in southern Siberia, is a well-known land-locked lake in the high latitude region. Research efforts have focused on its unique ecosystem and the stable stratification that historically defined it (*16–18*). However, a detailed analysis of long-term data on Lake Shira’s vertical structure reveals a significant shift in its mixing regime (*11*). During the ice-covered periods of 2015 and 2016, the lake underwent complete mixing. This event caused the temporary erasing its chemocline and the disappearance of hydrogen sulfide from the water column, signaling a transition from meromictic to holomictic conditions. This marked the first recorded instance of full mixing from 2002. It is hypothesized that strong wind currents associated with early ice melt during spring 2014 contributes to the lake’s complete mixing event in 2015 winter (*11*). The 2016 mixing appeared less pronounced, however, the position of mixolimnion was still deep to 23m (Bottom was 25m). By March 2017, mixing was shallow, and the lake reverted to meromictic conditions. These observations suggest that the lake’s mixing dynamics are highly variable and influenced by external environmental factors, such as wind, ice thickness, and freshwater inflows. These transitions also highlight how environmental perturbations can reshape microbial niches and redox structures even in systems previously considered physically stable.

Rogozin et al. (2017) study suggested that the mixing event in 2015 likely released nutrients from the monimolimnion into the photic zone, leading to an observed surge in organic carbon, chlorophyll a, copepod biomass (*Arctodiaptomus salinus*), and phytoflagellate (*Cryptomonas* spp.) abundance in early summer. Additionally, the breakdown of meromixis resulted in the disappearance of PSB from the monimolimnion, further highlighting the importance of investigating bacterial community composition changes and their environmental interactions during this period. Thus, this study focuses on such a critical rare event of change—when Lake Shira transitioned from meromictic to holomictic and returned to meromictic again—to investigate microbial responses across different mixing events. First, we investigated how microbial populations are distributed, their dynamics and adapt to the mixing events by applying 16S rRNA gene-based community composition at each water layer. Besides that, we also performed metagenomic sequencing to illustrated the microbes’ role in metabolic functions, nutrient cycling, and energy flow in different phases and layers (oxic and anoxic zones) of Lake Shira. We hypothesize that episodic holomixis disrupts the redox-driven vertical structuring of microbial communities, promoting the transient homogenization of metabolic niches, followed by rapid re-stratification upon the return of meromixis. We propose that these transitions reorganize both taxonomic composition and functional potential across depth layers. During mixing, the redistribution of reduced compounds such as sulfide, methane, and nitrogen intermediates may lead to the loss or relocation of specialist microbes like sulfur-cycling microbes, while enabling facultative anaerobes to expand their activity into upper layers. A set of microbial population contribute to key biogeochemical pathways and act as functional stabilizers of nutrient (sulfur, nitrogen, phosphate, etc.) cycling throughout dynamic regimes. Under favorable physical conditions, their cumulative activity may promote the concentration of reduced solutes, reinforcing the lake’s return to a stable meromictic state. This study provides a rare opportunity to capture the microbial and biogeochemical reassembly that follows the breakdown and restoration of stratification in a meromictic lake.

## Methods

### Sample Collection and the estimation of Limnological Parameters

Water sample were collected from Lake Shira (54.30’ N, 90.11’ E) which is located in Southern Siberia. We shared water samples with Rogozin et al. (*11*), who describe the detailed methods for sampling and limnological parameter collection.

A range of physicochemical parameters, including temperature, salinity, conductivity, redox potential, and dissolved oxygen, were measured in situ using either a YSI 6600 multiparameter probe (Yellow Springs, Ohio, USA) or a Hydrolab Data-Sonde 4a. After March 2015, dissolved oxygen concentration was inferred from redox potential due to equipment malfunction. Hydrogen sulfide levels were quantified by preserving samples with alkaline zinc carbonate and analyzing them using an iodometric titration technique(*19*). Conductivity measurements obtained in the field were normalized to a standard reference temperature of 25°C (K25) following modified ISO (1985) guidelines adapted for this specific lake (*8, 20*).

The water was collected from August 2015 to February 2019 at the deepest part of the lake (Suppl. Table S1). Fifty mL water samples were taken from various depths, store in -20°C until genomic DNA extraction. Total water sample number were 188. The “water-sediment interface” sample were collected from the upper surface soil when the lake in ice period of 2013, 2014, 2016, 2017, and 2018. The interface sample represents the boundary of water and sediment where the microbes sink within different lake hydrodynamic stage. To capture the microbial communities at the water-sediment interface with minimal disturbance, a custom-built sampler (freeze-corer made according to Renberg and Hansson (*21*) pre-filled with dry ice (CO_2_) and ethanol was used (Suppl. Fig. 1). When contact with the targeted interface, the sampler enabled freezing of the sample *in situ*, thereby preserving the structural and chemical integrity of the interface. The frozen samples were subsequently detached from the sampler, transported to the laboratory under cryogenic conditions. This procedure minimized anthropogenic interference during both collection and handling, ensuring the preservation of the interface microenvironment.

### Genomic DNA Extraction

Water sample were filtered on 0.2 μm 25 mm filter membrane (Whatman). And, the filter membranes were incubated in a solution containing 567 μL TE buffer, 30 μL of 10% sodium dodecyl sulfate (SDS), and 5 μL of RNase A (100 mg mL⁻¹) within 50 mL Falcon tubes. The incubation process was carried out at 37°C in a water bath for 60 minutes. Genomic DNA was subsequently extracted using the cetyltrimethylammonium bromide (CTAB) method, following established protocols. Genomic DNA products were used for both bacterial 16S rRNA gene amplification and metagenome.

### Bacterial 16S rRNA Gene Amplification

The bacterial 16S rRNA gene’s V6 – V8 hypervariable region was amplified from the extracted genomic DNA using primer pair U968F (5’-AACGCGAAGAACCTTAC-3’) and U1391R (5’-ACGGGCGGTGWGTRC-3’), generating a 424 bp fragment. The PCR reaction was performed in a total volume of 50 μL, containing 4 μL of 2.5 mM dNTP, 1 μL of each primer (10 μM), 0.5 μL of 5 U TaKaRa Ex Taq polymerase (Takara Bio, Otsu, Japan), 5 μL of 10× Ex Taq buffer, and 5 μL of template DNA (10–20 ng). PCR conditions included an initial denaturation step at 94°C for 5 minutes, followed by 30 cycles of denaturation at 94°C for 30 seconds, annealing at 52°C for 20 seconds, and elongation at 72°C for 30 seconds, with a final extension step at 72°C for 10 minutes. PCR products were separated on a 1.5% agarose gel and purified using the QIAEX II Agarose Gel Extraction Kit (QIAGEN, Hilden, Germany) according to the manufacturer’s instructions. The purity and concentration of the PCR products were assessed using a NanoDrop spectrophotometer (Thermo Fisher Scientific, Wilmington, Delaware, USA), while fragment length was confirmed via 1.5% agarose gel electrophoresis. Each purified PCR product was assigned a unique DNA tag through DNA-tagging PCR (DT-PCR). After DT-PCR, products underwent an additional purification step using the QIAEX II Agarose Gel Extraction Kit. Equal amounts (100 ng) of each purified DT-PCR product were pooled, and their concentrations were quantified using both a NanoDrop spectrophotometer and a Qubit fluorometer (Invitrogen, Carlsbad, CA, USA). The pooled library was then prepared for sequencing, and paired-end sequencing (2 × 300 bp) was performed using an Illumina MiSeq platform at Yourgene Bioscience (New Taipei City, Taiwan).

### Sequence Analysis

Raw paired-end reads were quality-checked with the QIIME2 (v2021.8) (*22*). Primer sequences were trimmed using cutadapt, and imported into QIIME2. Sequence denoising, merging, and chimera filtering were performed with the DADA2 plugin, using truncation lengths of ∼450 bp to ensure sufficient overlap for merging (≥20 bp). Amplicon sequence variants (ASVs) were identified directly from the filtered reads, and singleton features were excluded also by QIIME2. The ASVs were assigned taxonomy using the SILVA 138 database, with filtering to remove mitochondrial and chloroplast sequences. The final ASV table and representative sequences were exported for downstream diversity analysis and community composition profiling. The alpha (Chia and Shannon diversity) and beta diversity analyses were conducted using QIIME2 core-metrics-phylogenetic workflow. Samples were rarefied to a depth of 3,256 sequences per sample based on rarefaction curve saturation. Statistical comparisons of alpha diversity across environmental conditions or water layers were performed using non-parametric Kruskal–Wallis tests. Beta diversity was performed using both weighted and unweighted UniFrac distances as well as Bray–Curtis dissimilarity. Principal Coordinates Analysis (PCoA) was used to visualize dissimilarities in microbial community structure. Group differences in beta diversity were statistically evaluated using PERMANOVA via the “vegan” package in R (*23*). Taxonomic composition (relative abundance) was visualized using bar plots at phylum to genus levels.

To identify co-associations among microbes, genus-level co-occurrence networks analysis were constructed using relative abundance profiles. The analysis was performed in Rstudio 2024.09.0+375using the NetCoMi, psych, and tidyverse packages (*24–26*). For each stratification states, the genus abundance matrix was transposed and subjected to Pearson correlation analysis using psych package. Only the absolute correlation coefficient (|r|) ≥ 0.3. between genera were retained. For each pair, the correlation coefficient and *p*-value (<0.05) were extracted and compiled into a pairwise edge list. Networks were constructed as undirected graphs, where each node represented a genus and each edge represented a statistically supported co-occurrence by Gephi 0.10.1 (*27*). Topological patterns and potential keystone taxa were visualized and interpreted through Gephi 0.10.1 using the Circular layout, with edge width proportional to correlation strength and node size reflecting the relative abundance of each genus (*27*).

### Metagenomic library preparation, sequencing and analysis

To reveal the microbes’ functional profiles in distinct oxygenic condition and lake hydrodynamic stages, the metagenomic sequencing was performed on pooled samples from oxic or anoxic zone in each sampling period. That is, the genomic DNA was extracted from multiple depths within either the oxic or anoxic zones and pooled by four different lake hydrodynamic stages to generate a composite sample representing each condition. Pooling was performed due to each hydrodynamic stage comprised multiple sampling months and numerous sampling depth, making it impractical to use one or two depths as representative of the entire hydrodynamic stage. By pooling across depths and months, we aimed to capture the overall functional potential of the microbial communities while minimizing biases introduced by selecting individual depths. The DNA products from “interface” in 2016, and 2017, and 2018 were also performed metagenome. The DNA products quantified using the Qubit dsDNA BR assay kit in a Qubit 2.0 fluorometer. In total, eight metagenomic samples were prepared for whole-genome sequencing, including one sample represented “complete holomictic stage” (CH), one for the oxygen zone in “developing meromictic stage” (DM_OX), one for the anoxygen zone in “developing meromictic stage” (DM_AOX), one for the oxygen zone in “meromictic stage” (M_OX), one for the anoxygen zone in “developing meromictic stage” (M_AOX), and interface from 2016, 2017, and 2018. High-throughput long-read sequencing was carried out on a Nanopore PromethION platform at Biotools Inc. (Taipei, Taiwan). The generated Nanopore datasets were individually assembled using MetaFlye v2.9.3 (--nano-hq --meta) (*28*). To enhance consensus accuracy, the assembled contigs were iteratively polished with Racon (*29*) and subsequently refined with Medaka (https://github.com/nanoporetech/medaka). Raw sequencing data have been deposited in public repositories. The 16S rRNA gene amplicon reads are available in the NCBI Sequence Read Archive (SRA) under BioProject accession PRJNA1338226. Shotgun metagenomic sequencing reads are available in the European Nucleotide Archive (ENA) under BioProject accession PRJEB101372.

### Metagenome Assembly, phylogeny construction and functional annotation

To quantify the abundances of assembled contigs across individual samples, Nanopore reads were mapped to the assemblies using Minimap2 with default parameters (*30*). Contig abundance profiles were subsequently used by MetaBAT2 (*31*) to perform metagenome-assembled genome (MAG) binning. The quality of MAGs was evaluated with CheckM (*32*), and only those classified as medium quality (completeness ≥ 70%, contamination ≤ 10%) or high quality (completeness ≥ 90%, contamination ≤ 5%) according to the MIMAG standards (*33*) were retained for downstream analyses. Taxonomic classification and phylogenetic reconstruction of MAGs were carried out using GTDB-Tk v2.3.2 (*34*) against the GTDB reference database (Release 214) (*35*), applying default parameters (minimum alignment percentage = 10; genetic code = 11). The resulting phylogenies were visualized and annotated using the Interactive Tree of Life (iTOL) platform (*36*).

Protein-coding genes were predicted from both MAG-derived and unbinned contigs using Prokka (*37*), and functional annotation was performed with EggNOG-mapper v2.1.12, referencing the EggNOG 5.0.2 database (*38*). KEGG Orthology (KO) assignments derived from EggNOG were used to assess the metabolic potential of each MAG. Functional diversity was evaluated via principal coordinate analysis (PCoA) based on Bray–Curtis dissimilarity of KO profiles.

### CPM_contig_ value in four functional-related pathways

To quantify the relative abundance of genes involved in major biogeochemical processes (nitrogen, phosphorus, carbon, and sulfur cycles), we adopted the transcripts-per-million (TPM) normalization approach commonly applied in RNA-seq analyses (*39, 40*). Reads from each metagenomic sample were mapped to protein-coding genes on assembled contigs using Minimap2 (*30*) with default parameters. For each contig, read counts per open reading frame (ORF) and total mapped reads were extracted and used to calculate the contig-per-million mapped reads (CPM_contig_). CPM_contig_ values of ORFs associated with KEGG Orthology (KO) identifiers relevant to the four pathways were aggregated per sample to generate KO-level relative abundance matrices. These matrices were subsequently applied to functional profiling, including hierarchical clustering, Z-score–transformed heatmap visualization, and comparative analyses of gene expression patterns under oxic versus anoxic conditions and across meromictic versus holomictic phases.

## Results

### Hydrological Characteristics in Lake Shira from 2015 to 2019

Vertical profiling of sulfur concentration, temperature and salinity from 2015 to 2019 revealed clear temporal and seasonal stratification dynamic in Lake Shira (Fig. 1, Suppl. Fig. 2). Thermocline and halocline were observed from spring through autumn (May–October), with a warm, low-salinity surface layer overlying a colder, more saline monimolimnion—typical of a meromictic condition. However, during March and January 2015 and 2016, the thermal and salinity gradients were notably diminished. And, the sulfide concentration was under detection in all water column, suggesting a temporary mixing event (Fig. 1A). The depth of the mixolimnion was as deep as 23m, and the concentration of sulfide in this period was below detection. In contrast, the mixed layer reached 16m in March 2017. Also, the sulfide concentration gradually increased from August 2016 to 2017 (0.22 to 15.2 mg/L). It indicated a recovery to a meromictic regime of Shira Lake again in 2017. Thus, we defined this time period as “development meromictic stage”.

**Figure 1.**
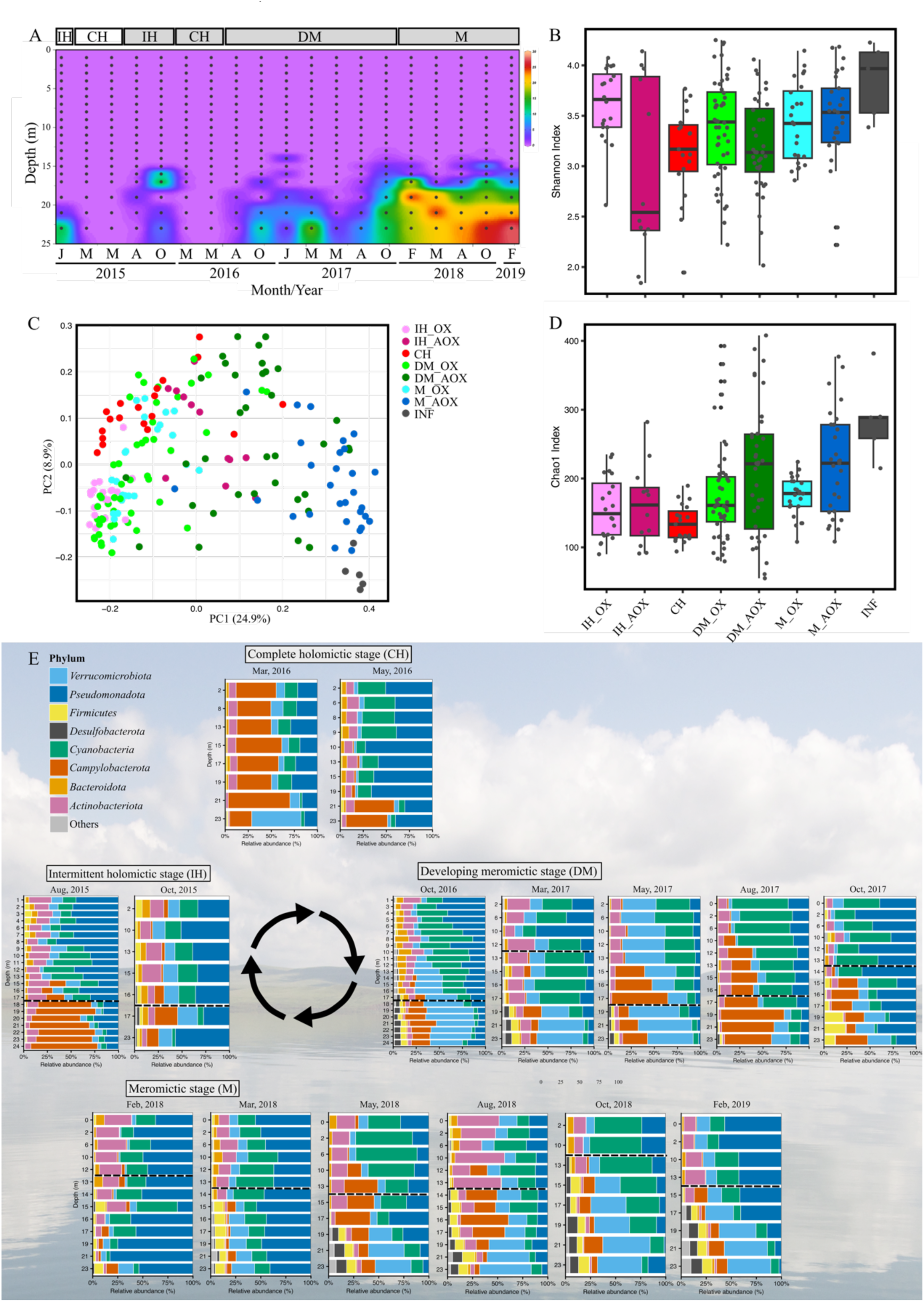
Environmental factor dynamics, microbial diversity, and stratification transitions in Lake Shira from 2015 to 2019. (A) Temporal variation of sulfide concentration across depths, showing episodes of intermittent holomictic stage (IH) in January and August–October 2015; complete holomictic stage (CH) in March–May 2015 and 2016; developing meromictic stage (DM) in August 2016 – October 2017; meromictic stage (M) in February 2018 – February 2019. The x-axis is labeled with month abbreviations, and years are indicated below. Gray boxes indicate time periods with 16S rRNA gene amplicon sequencing data. (B) Alpha diversity (Shannon index) of microbial communities across oxic (IH_OX, CH, DM_OX, M_OX), anoxic (IH_AOX, DM_AOX, M_AOX), and sediment–interface (INF) samples. (C) Principal coordinates of microbial community composition based on 16S rRNA gene amplicon data showing distinct clustering patterns for oxic (IH_OX, CH, DM_OX, M_OX), anoxic (IH_AOX, DM_AOX, M_AOX), and INF samples. (D) The transitions of vertical distribution of major bacterial phyla among four identified hydrodynamic stages—CH, DM, M, and IH. Bars represent the relative abundance of phylum-level bacteria groups by depth, illustrating the microbial community restructuring across time. Dashed lines on the depth axis indicate the approximate boundaries between oxic and anoxic zones.

After that, although the stratification in temperature and salinity profile exhibited less pronounced, the sulfide concentration remained high, reaching up to 25 mg/L at depths of 19 and 21 m during February and May 2018 (Fig. 1A). A strong thermal/salinity stratification was re-established after October 2018 (Supp. Fig. S2). In February 2019, profiles of both temperature and salinity remained stratified with a mixed layer extending to around 15 m. A high concentration of sulfide (28.13 mg/L at a depth of 23 m) was observed under the mixed layer, indicating the continued presence of a stable anoxic monimolimnion.

These hydrographic observations confirm that Lake Shira experienced periodic collapse and reformation of its vertical stratification, transition between the holomictic and meromictic stages. Based on sulfide profiles, we define the following stratification phases: intermittent holomictic stage (IH) in August–October 2015; complete holomictic stage (CH) in March–May 2016; developing meromictic stage (DM) in August 2016 – October 2017; meromictic stage (M) in February 2018 – February 2019. These stages provide a framework for interpreting microbial and biogeochemical dynamics presented in further investigations.

### Microbial Diversity Patterns Across Stratification Stages

A total of 4,845,913 high-quality sequences were obtained across all 193 samples after denoising and quality filtering. After removal of singletons and chimeras, 4,775 unique ASVs were retained. Rarefaction analysis indicated that all samples reached sufficient sequencing depth for downstream community analysis (Suppl. Fig. S3).

Bacterial community composition exhibited strong vertical stratification, corresponding closely to the changes of redox condition and lake mixing status. Also, the transition from holomictic to meromictic regimes was accompanied by marked shifts in community composition (Fig. 1D). Both the IH and CH stages featured relatively homogeneous communities throughout the water column, dominated by phylum *Pseudomonadota*, *Campylobacterota*, *Cyanobacteria*, and *Actinobacteriota*—reflecting well-mixed, oxic conditions. The *Cyanobacteria* were large accumulated in the sediment of INF2016 and INF2017 (Suppl. Fig. S4). Notably, *Campylobacterota* were broadly distributed across depths during CH but restricted to anoxic zones in the earlier IH stage, highlighting subtle redox-driven niche differentiation even during transitional phases (Fig. 1D). The homogenized conditions during CH likely limited niche differentiation, resulting in low taxonomic differences and reduced α-diversity variation, particularly evident in the Chao1 and Shannon profiles (Fig. 1B).

In contrast, the DM and M stages displayed strong vertical differentiation between oxic and anoxic zones (Fig. 1D). DM_OX waters were enriched in *Cyanobacteria* and *Pseudomonadota*, while anoxic bottom layers (DM_AOX) became increasingly dominated by *Desulfobacterota*, *Firmicutes*, and *Campylobacterota*, in line with the establishment of stable redox gradients. By M stages, community composition was clearly dichotomized in Class level (Fig. 2B and Suppl. Fig. 5). Anoxic bottom waters supported specialized anaerobic classes—including sulfate reducers and fermentative taxa (*Clostridia, Campylobacteria, Desulfobacteria*)—while surface waters remained dominated by phototrophic and aerobic heterotrophs (*Cyanobacteria, Alphaproteobacteria,* and *Actinobacteria*)(Suppl. Fig. 6). These patterns suggest the establishment of distinct microbial niches, maintained by strong and persistent chemical gradients. This period of partial stratification promoted increased community turnover, particularly between depth layers, as shown by the increasing Chao1 variability in anoxic bottom layers (DM_AOX and M_AOX)(Fig. 1B).

**Figure 2.**
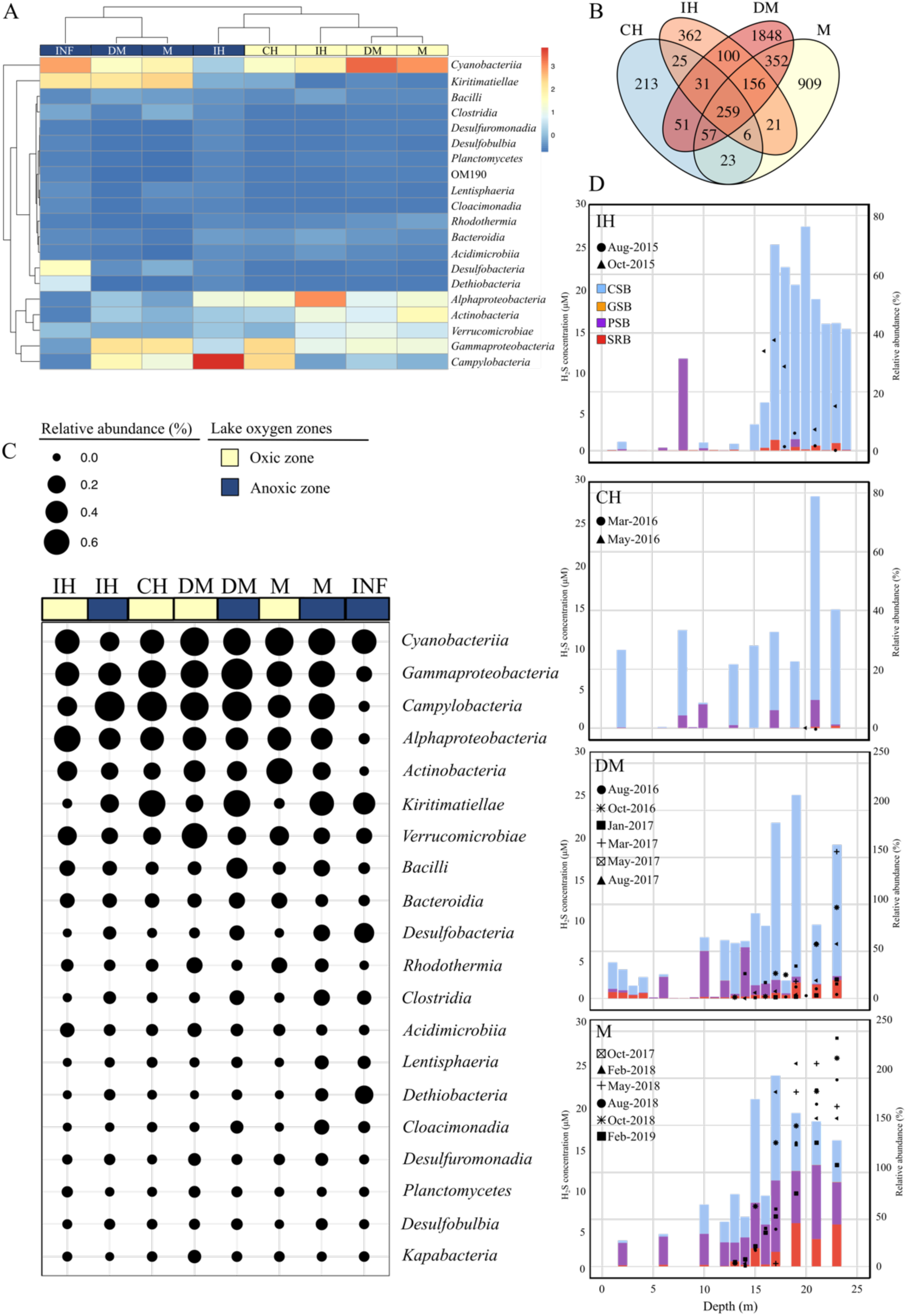
Spatiotemporal dynamics and stratification patterns of sulfur-cycling bacterial communities across lake mixing stages and redox zones. (A) Heatmap showing class-level relative abundance of major bacterial lineages associated with sulfur metabolism across sampling stages and oxygen zones. Samples are hierarchically clustered by Bray–Curtis dissimilarity. (B) Venn diagram depicting the number of unique and shared ASVs among the four defined stratification stages: intermittent holomictic (IH), complete holomictic (CH), developing meromictic (DM), and meromictic (M). DM and M stages harbored the highest numbers of unique ASVs, indicating increasing community specialization over time. (C) Dot plot summarizing the occurrence and relative abundance (circle size) of selected bacterial classes across oxygenation layers and stages. Color shading indicates oxic (light yellow) and anoxic (blue) zones. (D) Depth profiles of sulfur-metabolizing bacterial groups—including green sulfur bacteria (GSB), purple sulfur bacteria (PSB), colorless sulfur bacteria (CSB), and sulfate-reducing bacteria (SRB)—for each stratification stage (IH, CH, DM, M). Bars indicate relative abundance per depth; background blue shading represents measured sulfide concentrations (μM).

A PCoA analysis further illustrated the samples from oxic-layers (OX) clustered strongly and overlapped, PERMANOVA *p*=0.001), indicating that overall surface communities were compositionally consistent over time (Fig. 1C). The heatmap also supported by uniform patterns its observed which had dominance of *Cyanobacteriia*, *Alphaproteobacteria*, and *Actinobacteria* (Fig. 2A). The CH stage occupied a transitional position cross these community. The community in CH stage were classified as oxic condition; however, its samples were partially separated from the main OX cluster and located closer to IH_AOX from the PCoA analysis (Fig. 1C and Suppl. Fig. S7). This condition was consistent with the heatmap, where the CH samples clustered with IH_AOX and shared high relative abundance of *Campylobacteria*, *Alphaproteobacteria* and *Gammaproteobacteria* (Fig. 2A).

The community in anoxic samples (AOX) were distinct from that in OX group from the PCoA analysis. They showed broader distribution and greater dispersion across PCoA space, reflecting elevated beta diversity and compositional variability (Suppl. Fig. 7). However, compare with DM_AOX, the community in M_AOX samples were more uniform and appeared in the far-right quadrant of the plot, which closed with interface sediment (INF) samples. The anoxic layers, including INF, emerged in the deep layers with dominant of *Campylobacteria*, *Gammaproteobacteria*, and *Kiritimatiellae* (Fig. 2A). Furthermore, the dynamic of the community of INF from 2013 to 2018 showed that, after CH stage, a large accumulation of *Cyanobacteria* and *Kiritimatiellae* and reduced of *Desulfobacteria* in interface sediment in 2017 (INF_2017) (Suppl. Fig. S4 and S5). When the recovery of meromictic stage, the community in INF_2018 decreased of bacteria communities in *Cyanobacteria* and *Kiritimatiellae* and increased of the relative abundance of *Desulfobacteria*.

A Venn diagram of ASVs further illustrated compositional divergence across stages: a substantial number of stage-specific ASVs were detected in the M (n = 909) and DM (n = 1,848) stages, compared to fewer in IH (n=362) and CH (n=213) (Fig. 2C). In the core of Venn diagram, there were 259 ASV, which dominated by *Gammaproteobacteria*, *Actinobacteria*, and *Alphaproteobacteria* groups, constantly presented in every stage of stratification regime (Suppl. Fig. S8).

### Distribution of Sulfate-Reducing and Sulfur-Oxidizing Bacteria

The vertical distribution of key sulfur-metabolizing bacteria, including sulfate-reducing bacteria (SRB), colorless sulfur bacteria (CSB). purple sulfur bacteria (PSB), and green sulfur bacteria (GSB), varied significantly across stratification stages and depths (Fig. 2D and Suppl. Table S2).

During the IH stage, SRB—including members of *Desulfuromonadia* and *Desulfobacteria*—were present at low levels throughout the water column, with no clear depth stratification. Except PSB presented 30% of total bacterial community at a depth of 8 m (mainly *Rheinheimera* spp.), GSB and PSB were rare and undetectable, consistent with the low sulfide concentrations during this transiently mixed period (Fig. 2D). In contrast, CSB (mainly *Campylobacteria*) composited largely of bacteria community in the anoxic layers where the H_2_S concentration were 2-15 μM. In the CH stage, CSB with low relative abundance (20% to 80%) were more prevalent in every depth, while SRB remained largely absent or confined to low-abundance detections in deeper layers. These distributions mirrored the chemical profiles, where H₂S was not detected and redox gradients had yet to fully reform. PSB maintained low relative abundance from 0% to 9% in oxic to anoxic zone.

By the DM stage, CSB abundances increased markedly in anoxic layers. Simultaneously, SRB—particularly *Desulfobulbus*, *Desulfovibrio*, and

*Desulfomicrobium*—became enriched in bottom layers (18–25 m), coinciding with the accumulation of H₂S and clear thermal and salinity stratification. Notably, the PSB presented dominant near the oxic–anoxic interface while CSB and SRB were mainly in oxic or anoxic zones. In the anoxic layer of the lake, H₂S was accumulated again to 18 μM during this stage.

In the meromictic (M) stage, bacterial communities involved in sulfur metabolism became strongly stratified and functionally specialized along depth gradients. SRB (particularly *Desulfobacterium* and *Desulfoconvexum*) reached their highest relative abundances (exceeding 40%) in the deep anoxic layers (19-25 m), coinciding with H₂S concentrations >25 μM. These taxa dominated the sulfidic monimolimnion, actively mediating dissimilatory sulfate reduction. PSB (*Rheinheimera* spp.) also became substantially enriched during this stage. Unlike earlier stages, PSB were detected throughout the entire water column and were particularly abundant in the anoxic layers, where light and oxygen levels were low but sulfide concentrations were high. In addition, CSB-dominated by *Sulfurovum* spp. and *Sulfurimonass* spp.-were present in both oxic and anoxic layers, with peak abundance at the oxycline and suboxic interface. This tripartite spatial partitioning—with SRB dominating in the deep sulfidic zone, PSB broadly distributed but enriched in anoxia, and CSB concentrated near the redox interface—reflects a mature, functionally layered sulfur-cycling system in meromictic stage.

### Network Structure of Bacterial Communities Across Oxygenation Layers and different Stratification Stages

To find out potential microbial interactions and hub microbes, a co-occurrence network analysis revealed distinct patterns of microbial connectivity and modular organization across stratification stages and oxygenation layers (Fig. 3 and Suppl. Fig. S9). Each group-specific network (IH_OX, IH_AOX, CH, DM_OX, DM_AOX, M_OX, M_AOX, INF) was constructed based on genus-level relative abundance profiles using Pearson correlations, with node size representing relative abundance in total bacterial community and edge color indicating correlation direction which had significantly correlation.

**Figure 3.**
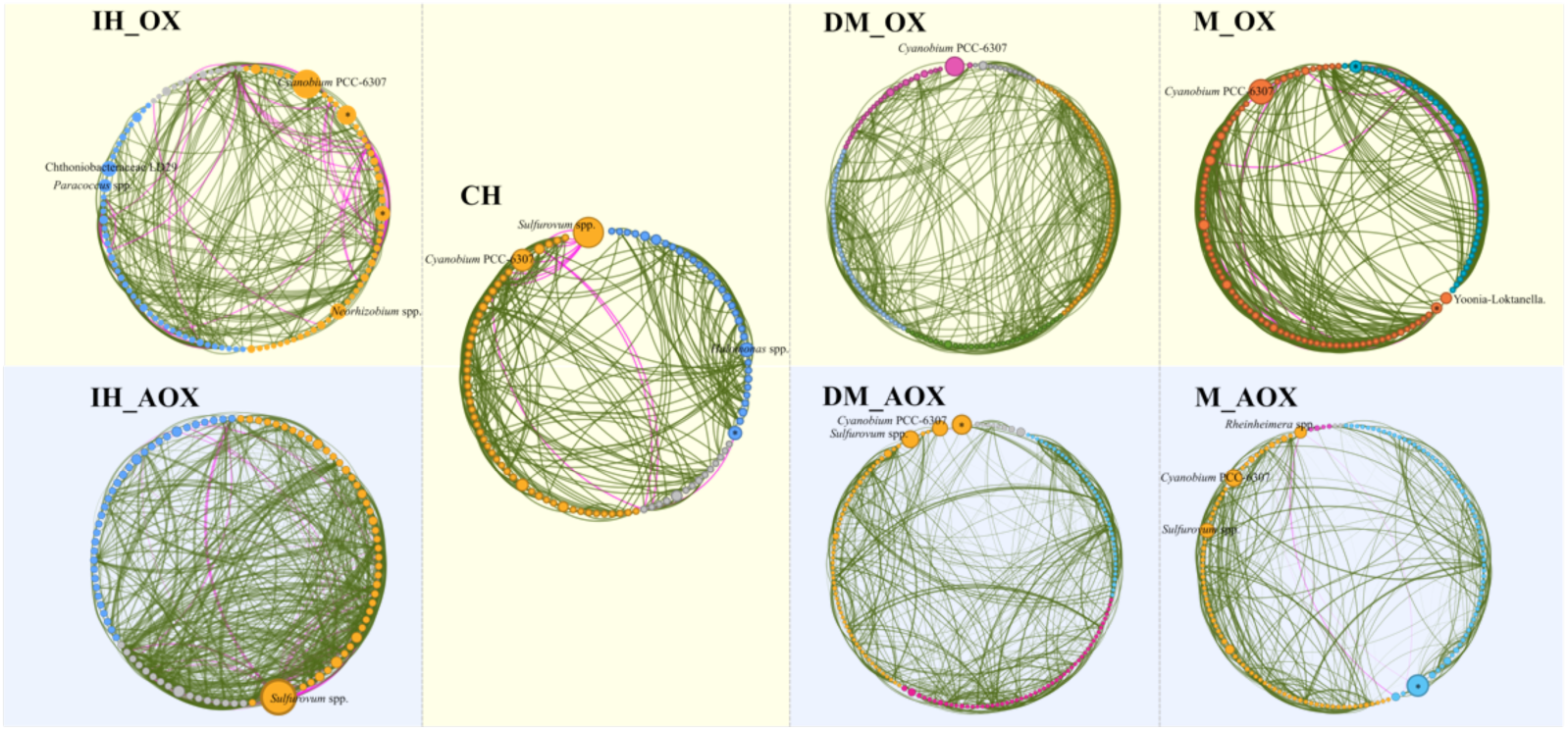
Co-occurrence networks of bacterial communities across different stratification stages and oxigenic zones in Lake Shira. Each circular network represents genus-level associations within a specific stratification stage and oxygenation zone: IH_OX, IH_AOX, CH, DM_OX, DM_AOX, M_OX, M_AOX. Nodes represent bacterial genera; node size corresponds to degree centrality (number of connections). Edges represent statistically significant pairwise Pearson correlations (|r| ≥ 0.3, p < 0.01), with green edges indicating positive correlations and magenta edges indicating negative correlations. Notable hub taxa are labeled, including *Cyanobium* PCC-6307, *Sulfuricurvum*…etc. *: unclassified bacterium. IH: intermittent holomictic; CH: complete holomictic stage; DM: developing meromictic stage; M: stable meromictic stage; INF: interface sediment. _OX: oxic zone; _AOX: anoxic zone.

During the IH_OX stage, most of associations were positive in the IH_OX network, although one distinct module (highlight in orange color) contained several negative association taxa. The IH_AOX network was denser overall, which show increased interconnections among taxa. Notably, *Sulfuricurvum* spp. was the major hub, which had both positive and negative correlations with many taxa. It suggesting early establishment of sulfur-metabolizing lineages even under transitional anoxic to oxic conditions. And, many genera retained similar relative abundance and positive interactions compared with the IH_OX network, reflecting a structural differentiation. In the CH stage, the network appeared more centralized, with *Sulfurovum* spp., *Cyanobium* PCC-6307, and *Halomonas* spp. remaining as highly connected nodes, particularly *Sulfurovum* spp. displayed negative associations with numerous taxa. And, thick positive and negative correlation lines in CH network presented highly association with each other. As the lake transitioned from complete mixing to the onset of persistent stratification during the DM stage, this phase represents the first fully expressed divergence between surface and deep-water microbial communities following the holomictic period. In the DM_OX network, overall connectivity increased compared to earlier oxic networks, marked by a larger number of positive correlations among low-abundance taxa. However, the network was dominated by a single highly abundant node, *Cyanobium* PCC-6307, while most other genera remained at relatively low abundance. This pattern reflects a community still shaped by surface oxygen availability but beginning to reorganize as vertical gradients strengthen. In contrast, the DM_AOX network exhibited higher taxonomic richness and greater relative abundance among multiple taxa, including *Sulfurovum* spp., and *Cyanobium* PCC-6307. The increased abundance and diversity of central nodes in DM_AOX indicate a more functionally differentiated and ecologically structured community, adapted to emerging anoxic and sulfidic conditions. Notably, both DM_OX and DM_AOX networks exhibited more numbers of distinct co-occurrence modules (partitions) than observed in other stages, suggesting increased modularity and niche segregation as the water column became chemically stratified. This increase in network complexity reflects the establishment of redox-driven ecological boundaries and the initial emergence of sulfur-metabolizing community frameworks that dominate in the beginning of the meromictic stage.

The M_OX network was characterized by a large number of strong positive correlations, forming a dense connectivity web among taxa. Despite this elevated overall connectivity, the network contained only two distinct co-occurrence modules (partitions), suggesting relatively low community modularity. The dominant taxa included *Cyanobium* PCC-6307—consistently abundant across stages—and a newly node, *Yoonia-Loktanella*, which contributed high relative abundance in this network. These patterns indicate that, M_OX communities maintained tight internal associations with relatively compositionally simple in this network (Suppl. Fig. S10). In contrast, the M_AOX network exhibited higher modularity, comprising four distinct co-occurrence modules, but showed fewer and weaker correlations overall. Key taxa within this network included *Sulfurovum* spp., *Cyanobium* PCC-6307, and *Rheinheimera* spp., each acting as abundant nodes within this network. The presence of these genera, coupled with limited cross-module connectivity, suggests distinct ecological clusters occupying specialized niches within the anoxic zone.

In final, the INF (interface sediment) network exhibited a uniquely complex structure compared to all stages (Suppl. Fig. S9). It displayed strong/highly positive correlations as well as a substantial number of strong negative associations, distinguishing it as the most interaction-rich across all stratification phases. While *Cyanobium* PCC-6307 remained a consistently abundant and highly connected node, the majority of dominant taxa in this network were unclassified at the genus level, highlighting the ecological distinctiveness and limited characterization of benthic microbial assemblages.

### Metagenome analysis in different oxygenation layers and Stratification Stages

Metagenomic sequencing was performed on five composite samples (CH, DM_OX, DM_AOX, M_OX, M_AOX), representing pooled oxic and anoxic layers from each stage to complement the amplicon-based taxonomic profiles. Two metagenome data from little-disturbed water-sediment interface in ice period of 2016, 2017 and 2018 (INF_2016, INF_2017, and INF_2018) which represented the time before CH stage and the time before M stage. Following metagenome assembly and quality filtering, 401 metagenome-assembled genomes (MAGs) were recovered. Of these, 268 MAGs were of medium quality (completeness ≥ 50%, contamination ≤ 10%) and 133 were of high quality (completeness ≥ 90%, contamination ≤ 5%) (Suppl. Table S3). Taxonomic classification assigned 399 MAGs to Bacteria and 2 to Archaea, encompassing nine distinct prokaryotic phyla (Fig. 4A; Suppl. Table 4). Bacterial MAGs spanned 21 phyla, with five dominant groups: *Pseudomonadota* (108 genomes, 26.93%), *Actinomycetota* (83 genomes, 20.70%) *Bacteroidota* (49 genomes, 12.22%), *Verrucomicrobiota* (40 genomes, 9.98%) and *Planctomycetota* (33 genomes, 8.23%) (Fig. 4A and Suppl. Table S5 to S8). Several rare MAGs from less-characterized phyla, such as *Gemmatimonadota* and *Cloacimonadota* were also recovered, particularly from anoxic samples (DM_AOX and M_AOX), suggesting the presence of underexplored anaerobic functional guilds in these environments (Fig. 4A).

**Figure 4.**
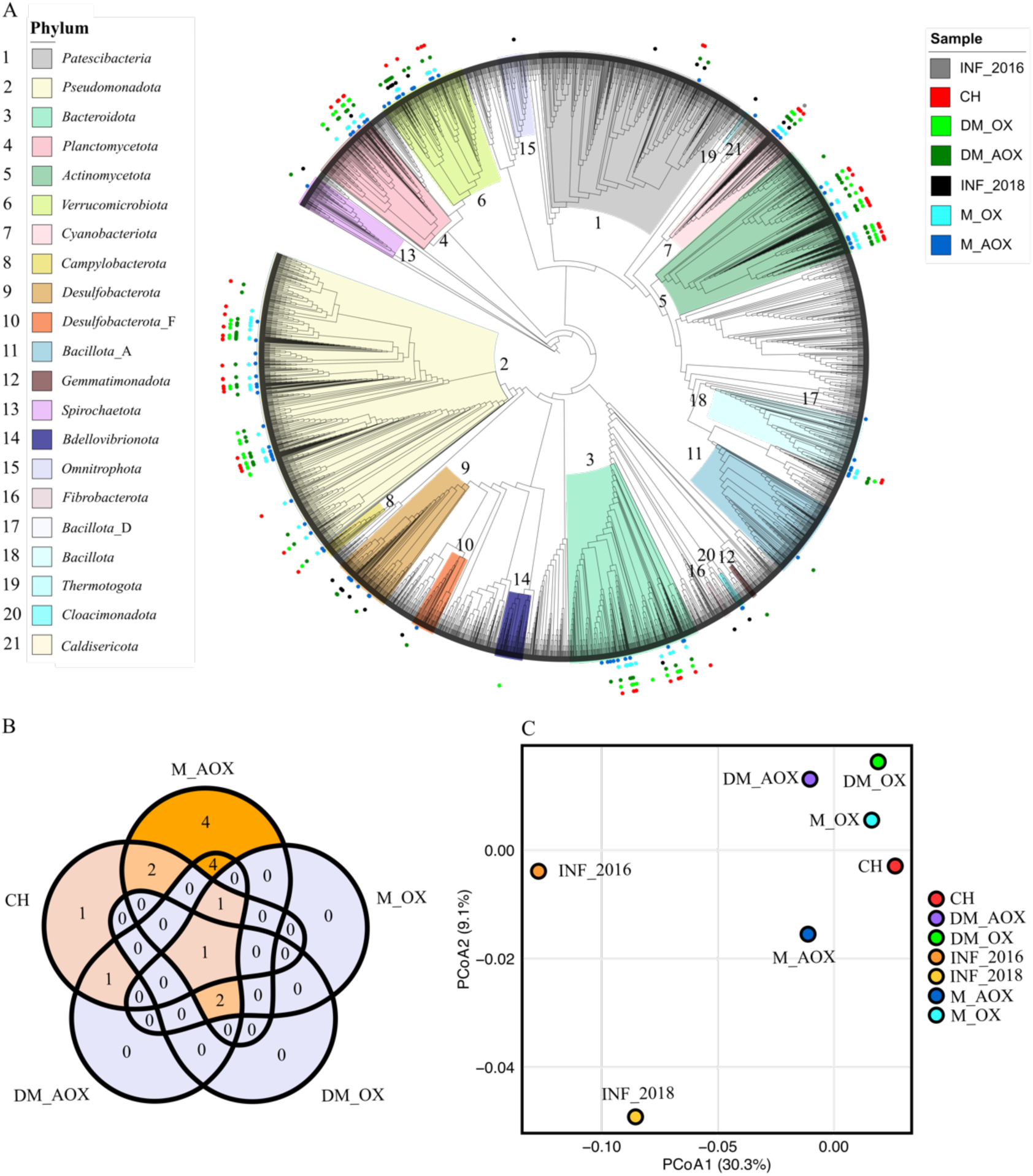
Taxonomic and compositional overview of metagenome-assembled genomes (MAGs) recovered from Lake Shira. (A) Phylogenomic tree of 399 dereplicated MAGs constructed. Branches are colored by phylum-level taxonomy (see legend at top left); sample origin is indicated by colored dots on the tree tips (see legend at top right). (B) Venn diagram showing the number of shared and unique sulfur-related MAGs across different lake phases and oxygenation zone. (C) NMDS ordination based on Bray–Curtis dissimilarity of KO composition across samples, illustrating functional differences among MAG communities, colored by sample. CH: complete holomictic stage; DM: developing meromictic stage; M: stable meromictic stage; INF: interface sediment. _OX: oxic zone; _AOX: anoxic zone.

A Venn diagram showed that 21 bacterial phyla, including 5 *Bacteroidota*, 8 *Actinomycetota*, 1 *Cyanobacteriota*, 2 *Planctomycetota*, 4 *Pseudomonadota*, 1 *Bacillota* (Suppl. Fig. 11A and Suppl. Table S9). Specify, when considering the anoxic (AOX) environments, the number of stage-specific phyla detected only in M_AOX was higher than those in DM_AOX, suggesting a more distinct microbial assemblage associated with the fully developed meromictic condition. For oxic layers, the number of stage-specific phyla in DM_OX (n = 19) also exceeded those detected in CH and M_OX stages. Furthermore, the sulfur-related MAGs were recovered 11 MAGs and 15 MAGs in DM_AOX and M_AOX stages (Suppl. Table S10 and S11), including many distinct SRB, PSB, PNSB, CSB, and SOB groups. Among them, there was one core sulfur-related MAG, which was the *Yoonia* spp. (*Pseudomonadota, Alphaproteobacteria, Rhodobacterales, Rhodobacteraceae*), was revered from all stages (Fig. 4B). The relative abundance of this PNSB (*Yoonia* spp.) were especially high in CH and M_OX stages (Suppl. Fig. 10).

### Functional Differentiation of MAGs Based on KO Profiles

The KO profiles of recovered MAGs from different stratification stages and hydrodynamic layers represented the overall functional divergence among microbial populations. The presence or absence of specific functional pathways across MAGs, indicating the functional potential of locally adapted populations. The barplot summarizes the proportion of MAGs carrying genes associated with specific metabolic pathways, grouped by different stratification stages. MAGs from DM_AOX and M_AOX samples carried higher proportions of genes related to sulfur metabolism, stress response, and phosphorus metabolism (Suppl. Fig. 11B).

A NMDS ordination based on MAG KO profiles revealed a clear separation of functional gene content among stratification stages and oxygenation zones (Fig. 4C). MAGs derived from CH, DM_OX, and M_OX samples clustered tightly together, indicating that genomes adapted to oxic surface layers and shared a similar set of metabolic traits. In contrast, MAGs from M_AOX and INF samples closed with each other, positioned far from the oxic group. This reflects the functional convergence of genomes adapted to stable anoxic and sulfidic environments, enriched in genes for sulfur respiration, fermentation, and methanogenesis. Importantly, M_AOX MAGs were positioned closest to the same-year INF samples (e.g., M_AOX and INF_2018), suggesting that microbial communities in the deep anoxic water column and benthic interface.

The NMDS distribution also mirrors the temporal progression of stratification. MAGs from CH—representing the fully mixed holomictic phase—aligned with oxic-layer MAGs (DM_OX and M_OX) due to shared access to light and oxygen (Fig. 4C). As the lake transitioned into the DM stage, functional profiles began to shift: DM_OX and DM_AOX MAGs remained close to oxic-layer MAGs. But, the DM_AOX MAGs started diverging toward an anoxic functional signature. By the M stage, with stratification fully established, M_AOX MAGs acquired KO compositions highly similar to those found in sediment-derived MAGs from INF samples.

To assess how microbial functional potential varies across all stages, we profiled the distribution of 173 key KEGG Orthologs (KOs) across 375 MAGs recovered from five stratification stages. MAGs from the oxic layers (CH, DM_OX, and M_OX) consistently harbored genes involved in aerobic sulfur oxidation, including components of the Sox multienzyme system (*soxA/B/C/X/Y/Z*) and the sulfite-oxidizing complex *soeB*/*C* (Suppl. Fig. S12). These MAGs exhibit a widespread capacity for sulfite oxidation as an energy-conserving process, which a sulfur-oxidizing bacteria under oxygenated conditions. Phototrophic potential was also prominent among oxic MAGs. The presence of carotenoid biosynthesis genes (*crt* cluster), bacteriochlorophyll synthesis genes (*bch* cluster), and light-harvesting complex proteins (*pufA*, *pufL*, *pufM*), suggests the presence of photoheterotrophic or anoxygenic phototrophic populations, likely active in surface waters or localized microaerobic niches such as organic aggregates or biofilm-like structures. Notably, several MAGs affiliated with *Planctomycetota* encoded methanogenesis-related genes, including *cdhE, ftr,* and the *fwdA/B/C*, across all oxic stages (CH, DM_OX, M_OX). MAGs from anoxic layers had the enrichment of *Desulfobacterota* MAGs, which carried a suite of methanogenesis genes, including *cdhC*, *cdhD*, *cdhE*, and the *fwdA–F* gene cluster. Also, these same MAGs carried marker genes of dissimilatory sulfate reduction, including *qmoA/B/C*, tus-dsrH/F/E, and *mvhD*, as well as nitrogen fixation genes (*nifH/D/K*) and nitrate reduction pathways such as *napA/B*, *narG/H/I*, and *nirK/S*. The DM_AOX layer hosted an especially diverse assemblage of *Pseudomonadota* MAGs that encoded both sulfate oxidation and reduction genes—including *soxX/Y/Z*, *soeB/C*, *hdrA*/B, and *aprA/B*—alongside nitrogen fixation (*nifH/D/K*) and nitrate reduction genes (*narG/H/I*, *nirB*, *napA/B*). Across all five stages, genes involved in nitrate/nitrite transport (*nrtA*–*D*, *NRT2*) and ammonia assimilation or release (*glnA*, *gltB*, GLUD1_2, *arcC*) were broadly distributed in all MAGs.

A total of 21 core MAGs were identified, and the presence–absence profiles of 48 KOs revealed conserved taxonomy and their metabolic lists (Fig. 5). These core MAGs, spanning 11 bacterial classes in *Alphaproteobacteria*, *Actinomycetia*, *Gammaproteobacteria*, *Bacteroidia*, and one phylum *Planctomycetia*, encode a stable set of genes that collectively support sulfur, nitrogen, methane and carbon cycling across all stages. Sulfite oxidation potential was widespread among nearly all core MAGs, with genes such as *soxB*, *soxC*, *soxX*, and *soxY* present across multiple taxonomic groups. Also, genes involved in central carbon metabolism, including the citrate cycle (*sdhB*, *mdh*, IDH3) and fatty acid degradation (ALDH), were broadly distributed. Among the core set, a distinctive subgroup of five MAGs—including two *Alphaproteobacteria*, one *Gammaproteobacteria*, one *Cyanobacteria*, and one *Planctomycetia*—stood out for their unusually broad functional scope. These MAGs carried genes associated with all major nutrient cycling pathways, enclosed sulfur metabolism, nitrogen fixation and respiration, methanogenesis, as well as phototrophic functions, including phycobilisome assembly (*cpcA/B/G*) and Calvin cycle components (*rbcL*, PRK). In addition, several core MAGs (*Actinomycetia*) consistently encoded a subset of key sulfur oxidation genes. Four MAGs belonging to the phylum *Bacteroidota*—classified within the *Bacteroidia* and *Rhodothermia* classes—carried genes associated with both sulfur oxidation and reduction pathways, as well as the nitrogen fixation operon (*nifH/D/K*).

**Figure 5.**
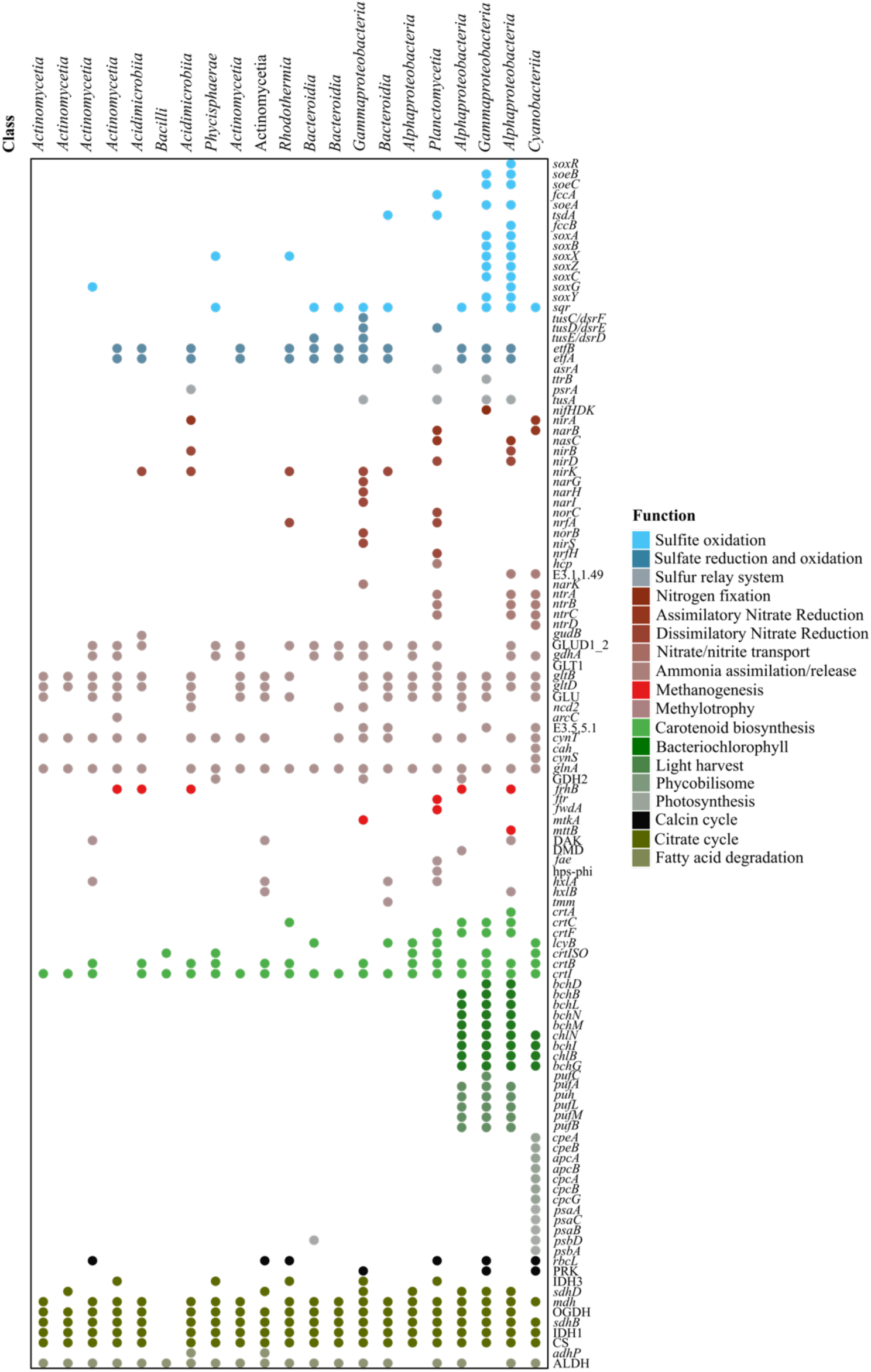
Functional gene profiles of 21 core MAGs recovered across all lake stratification stages. Presence–absence heatmap of 48 key KEGG Orthologs (KOs) detected in 21 metagenome-assembled genomes (MAGs) that were consistently found in all five hydrodynamic stages. Columns represent individual MAGs class name, and rows correspond to KOs grouped by functional category (right).

### Functional Genes and Sulfur, Nitrogen, Carbon, and Phosphate Metabolism Pathways

The abundance of key metabolic genes involved in sulfur, nitrogen, and carbon cycling using CPM-normalized values derived from metagenomic reads (Fig. 6 and Suppl. Fig. S14). The heatmap revealed distinct biogeochemical signatures in oxic vs. anoxic zones and across hydrodynamic stages.

**Figure 6.**
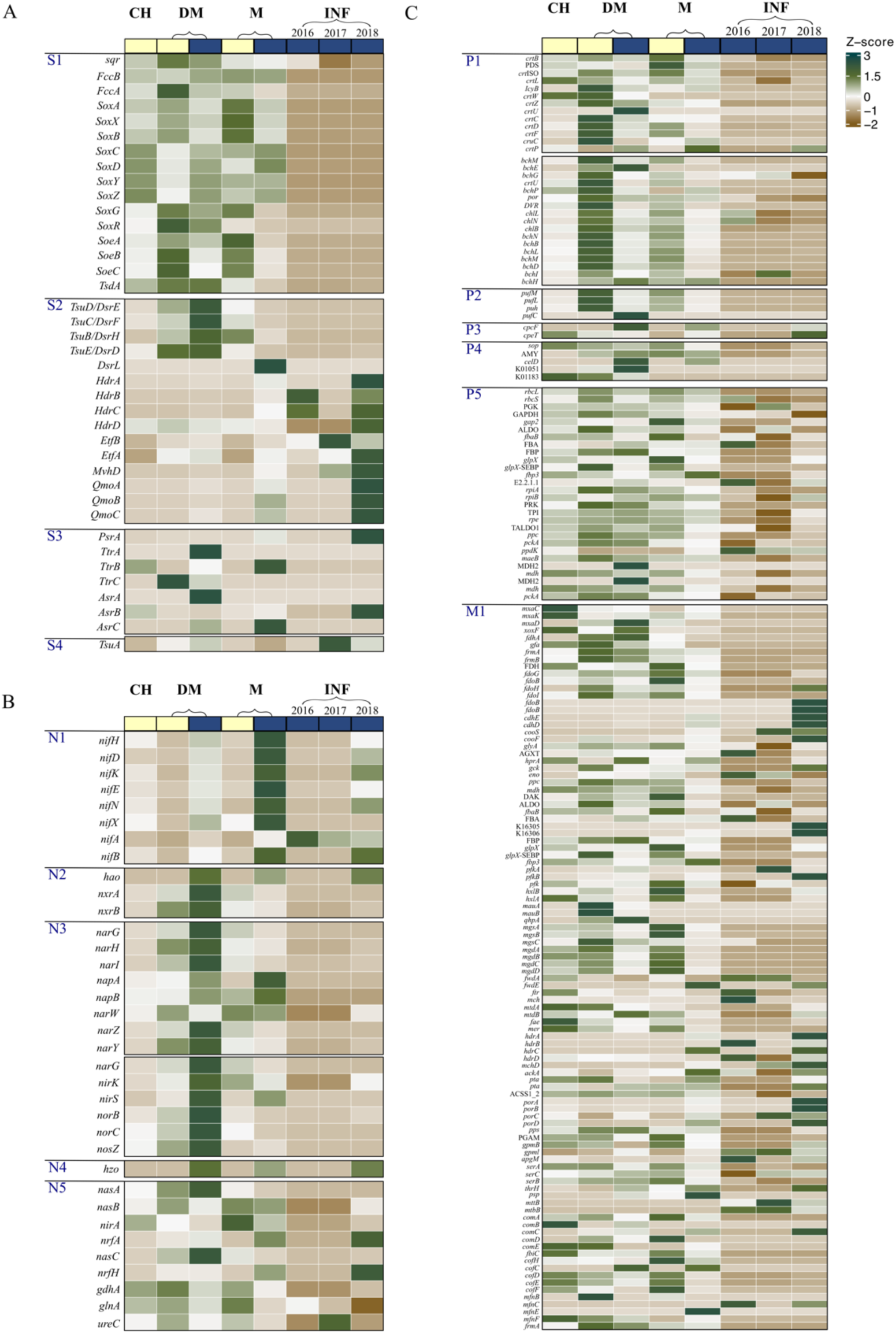
The distribution of key metabolic genes involved in sulfur, nitrogen, methane, and carbon cycling across lake stratification stages and interface sediment (INF). Heatmap displays Z-score normalized CPM abundances of representative KEGG Orthologs (KOs) associated with major biogeochemical processes, calculated from metagenomic reads across five stages and oxygenic layers. Genes are grouped into four functional groups: (1) sulfur metabolism (S1–S4): S1, sulfur oxidation; S2, dissimilatory sulfate reduction / Dsr-associated sulfur oxidation; S3, sulfur-cycle intermediate metabolite reduction; S4, sulfur relay system, (2) nitrogen metabolism (N1–N5): N1, nitrogen fixation; N2, nitrification; N3, denitrification; N4, anammox; N5, nitrate reduction and ammonification (DNRA-related processes), (3) Phototrophy/carbon cycling and methane cycling modules (P1–P5, M1): P1, enzymes of bacteriochlorophyll and carotenoid biosynthesis; P2, assembly of photosynthetic complexes; P3, phycobilisome; P4, non-chlorophyll phototrophy decomposition; P5, Calvin cycle; M1, methane oxidation and methanogenesis. CH: complete holomictic stage; DM: developing meromictic stage; M: stable meromictic stage; INF: interface sediment. Light yellow box: oxic zone; Blue box: anoxic zone.

### Sulfur metabolism

Genes involved in sulfur oxidation (e.g., *soxA/B/X/Y/Z*, *sqr*, *fccA/B*) were broadly distributed across samples, but reached highest TPM values in oxic and suboxic samples, particularly in DM_OX and M_OX (Fig. 6A). In contrast, genes involved in sulfate reduction, such as *dsrE/F*, *hdrA/B/C/D*, and *qmoA/B/C*, were concentrated in anoxic metagenomes, especially in DM_AOX, M_AOX and INF. Many sulfur reduction reactions increased especially in INF_2018. Also, two genes linked to intermediate sulfur transformations (e.g., *ttrA/B* and *asrA/C*) were enriched in both INF and deep anoxic layers (DM_AOX and M_AOX). In contrast, during the CH stage, sulfur metabolic gene expression was minimal across all sulfur oxidation, reduction, and intermediate transformation pathways. TPM values of key genes such as *soxB*, *sqr*, *dsrE/F/H/D*, and *ttrA/B* were uniformly low, indicating a suppression of active sulfur cycling processes.

### Nitrogen Metabolism

Nitrogen cycling genes showed stage- and layer-specific patterns. Genes for nitrogen fixation (*nifH/D/K/E/N/B*) were abundant in anoxic layers, especially in M_AOX (Fig. 6B). Both Nitrification genes (*nxrA/B*, *hao*), denitrification genes (*nirK*, *norB*, *nosZ*), and anammox (*hzo*) were peaked in anoxic layers (e.g., DM_AOX, M_AOX), reflecting canonical anaerobic denitrification processes. Interestingly, genes related to dissimilatory nitrate reduction to ammonium (DNRA) and ammonification were observed in both interface (INF_2018) and anoxic layers, suggesting alternative nitrogen retention strategies under low-oxygen conditions. During CH stage, nitrogen metabolism was dominated by assimilatory pathways. High TPM expression of *nirA*, *gdhA*, and *glnA* indicated active nitrate reduction to ammonium and its incorporation into amino acids. These patterns suggest that, under well-oxygenated and mixed water column conditions, microbial communities primarily engaged in nitrogen assimilation rather than dissimilatory or anaerobic nitrogen processes.

### Carbon Fixation and Phototrophic Metabolism

Genes involved in carbon fixation pathways were detected, including the Calvin cycle (*rbcL*, *prkA*), pigment biosynthesis (bacteriochlorophyll and carotenoid), assembly of photosynthetic light-harvesting complexes and phycobilisome (Fig. 6C).

Genes associated with bacteriochlorophyll and carotenoid biosynthesis (e.g., *bchN/B/L/M/D/I/H*, *crtB/W/Z/U/C/D/F*, *crtL/W/Z*, *IcyB*) photosystem assembly (*pufM/L/C*) and phycobilisome components were detected mainly in DM_OX and M_OX while these gene were rather decreased in CH stage. Also, these genes were present at lower levels in anoxic metagenomes (DM_AOX and M_AOX). Calvin cycle genes (e.g., *rbcL*, *prk*, *gapA*) were most abundant in oxic layers (especially in M_OX and DM_OX) while they were relatively low especially in the sample from INF (INF_2016, INF_2017, and INF_2018).

### Methane Oxidation and Methanogenesis

Most methane oxidation and methanogenesis genes were more highly detection in oxic layers (DM_OX and M_OX) and during the CH stage, consistent with well-oxygenated conditions supporting aerobic methanotrophy (Fig. 6C). Key genes associated with methane oxidation—including *mxaC/K/D*, and *xoxF*—were particularly enriched in CH and M_OX samples. These genes encode particulate and soluble methane monooxygenases and methanol dehydrogenases, typical of aerobic methanotrophs. However, several methane-related genes also showed notable enrichment in anoxic layers and INF samples, suggesting the presence of anaerobic or facultatively anaerobic methane-processing taxa. For example, *mxaD*, *cdhE/D*, *porA/B/C*, and *fdoB* elevated in M_AOX and INF, with the highest detection frequency observed in INF_2018, pointing to potential anaerobic methylotrophy or redox-linked methane utilization in these deeper, anoxic habitats.

### Citrate Cycle, Fatty Acid Degradation, Phosphate Related Metabolic Pathways

Genes involved in the central carbon metabolism and energy generation, such as *scuD/C*, *frdA/C/D*, *mqo*, CS, DLD, were widely distributed across samples but were particularly enriched in DM stage (Suppl. Fig. S14). In contrast, TCA cycle gene expression decreased in M_AOX and INF2016. Genes associated with fatty acid degradation, including *fadA/B/E* and *adhE/P/Y*, showed broad detection across all stages, but exhibited notably higher expression in DM stage, particularly DM_OX.

Genes associated with phosphate metabolism, including phosphate and phosphonate metabolism, phosphate related transporters, and pentose phosphate pathway. Basically, most of phosphate-related genes were broadly detected across all samples but were particularly enriched in oxic layers during DM and M stage. INF_2018 had relative high detection of phosphate-related genes rather than in INF_2016 and INF_2017.

## Discussion

In meromitic lake’s stratification maintained by salinity, light, and temperature, distribution of oxygen, sulfide, and inorganic nitrogen establish redox gradients that govern access to energy and nutrients, shaping the microbial community composition, dynamics and interactions. When any alteration, disturbances or even holomixis, chemical fluxes and energy gradients are reset, triggering relays of restructuring in communities and metabolism. Yet, in meromictic systems, mixing is an uncommon and unpredictable; coupled with the practical difficulty of obtaining samples in ice cover season and the water-soil interface, constraining sustained, continuous observations of microbial succession. Here, we integrate high-resolution sequential samples with amplicon and metagenomic data in four states of Lake Shira-intermittent holomictic (IH), complete holomictic (CH), developing meromictic (DM), to meromictic stage (M) of Lake Shira. And, we also characterize microbial community shifts and potential metabolic functions across these transitions, providing insights into how microbial assemblages reorganize and adapt to rare but ecologically significant disruptions in lake stratification (Fig. 7). To the best of our knowledge, this represents the first study to characterize microbial–environmental temporal dynamics during a rare holomixing episode in a long-term stable meromictic lake. The following will discuss dynamics of microbial communities and their potential metabolic networks at 4 different stages of the lake structures.

**Figure. 7.**
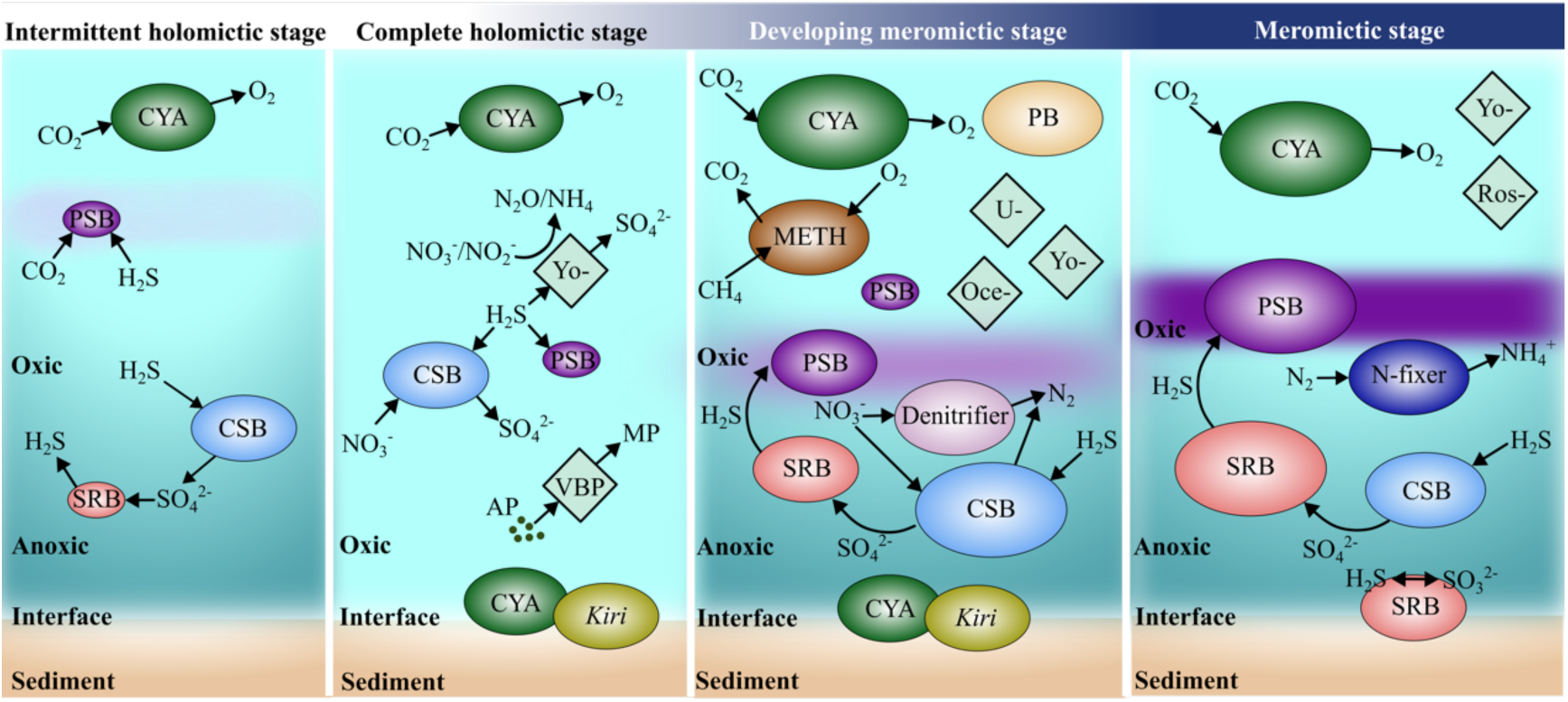
Schematic Illustration of redox-partitioned microbial trophic structure and metabolite exchange across hydrodynamic stages in Lake Shira. The schematic summarizes the dominant microbial taxa inferred for each stage—intermittent holomictic (IH), complete holomictic (CH), developing meromictic (DM), and meromictic (M)—and their putative metabolic linkages across the oxic layer, anoxic layer, and water–sediment interface. Arrows denote predominant transformation pathways and exchange of key substrates/products (CO₂, O₂, H₂S, SO₄²⁻, NO₃⁻/NO₂⁻, N₂/NH₄⁺, CH₄), highlighting stage-dependent coupling among oxygenic primary production, phototrophic and chemotrophic sulfur oxidation (PSB/CSB), sulfate reduction (SRB), nitrogen loss and recycling (denitrifiers; N-fixers), and methane cycling (METH). Labeled nodes represent major guilds and representative taxa inferred to contribute to these processes: cyanobacteria (CYA), purple sulfur bacteria (PSB), colorless sulfur bacteria (CSB), sulfate-reducing bacteria (SRB) from 16S rRNA analysis. Other abbreviations, Methylotrophs (METH), bacteriochlorophyll- and carotenoid-producing bacteria (PB), denitrifiers, nitrogen fixers (N-fixer), *Kiritimatiellae* (Kiri), *Yoonia* (Yo-), *Roseicyclus* (Ros-), *Oceanicaulis* (Oce-), UBA2463 sp018402635 (family *Burkholderiaceae*; U-), and a composite group of *Verrucomicrobiota*, *Planctomycetota*, and *Bacteroidota* (VBP) were the representative MAGs. Putative organic-matter sources and intermediates are indicated as algal/gel-like sulfated polysaccharides (AP) and monomeric polysaccharides (MP). The scheme highlights how holomixing resets chemical gradients and redistributes functional potential, whereas restratification re-establishes vertically partitioned niches that intensify redox-coupled sulfur–nitrogen–carbon interactions.

### Intermittent holomictic stage

The transition from IH to CH in Lake Shira induced a pronounced reconfiguration of microbial communities, particularly in the monimolimniion layers. A complete mixing event occurred in spring 2015. Although the lake appeared to progress toward a develop in meromictic state by August and October 2015, unusually thin ice cover in winter 2015 weakened salinity gradients, and the spring 2016 mixing depth remained deep. Thus, the system returned to holomictic conditions in March and May 2016. The IH period therefore represented a short transition phase situated between two CH states: an anoxic layer emerged but quickly disappeared as the mixed layer deepened, driving rapid shifts in physical and chemical conditions. The anoxic layer in IH stage shows a high proportion of positive and negative associations with tightly knit modules, indicating syntrophy-driven assemblages shaped by strong variance environmental filtering under reducing conditions and co-variation in response to shared resources. Concomitantly, community evenness increased, reflecting coordinated increases or declines across multiple taxa during this rapid reorganization. Taxa capable of tolerating and exploiting microaerophilic, sulfidic niches became prominent, including *Campylobacteria* (genus *Sulfurovum*) and *Alphaproteobacteria* (genus *Paracoccus* and SAR11 clade III). These groups frequently reported from hydrothermal/cold seeps, sulfidic caves and sediments, and redoxclines(*41, 42*). Although there is no *Paracoccus* MAG was recovered in this study, their known capacities for nitrate reduction and complete denitrification under low-oxygen conditions from previous studies (*43, 44*). Four genus *Sulfurovum* MAGs (class *Campylobacteria*, former class *Epsilonproteobacteria*), from this study, were detective in both oxic and anoxic layers of DM and M stage. Genus *Sulfurovum* had genetic capability to oxidize sulfide, thiosulfate and sulfite, and to reduce polysulfide. And, these organisms appear genetically adapted to thrive in the increasing sulfide concentrations and low-oxygen niches characteristic of lake waters (*45, 46*). Moreover, subtle variations in genetic potential—especially those related to redox processes and oxygen tolerance—likely facilitate fine-scale niche partitioning among lake populations (*47, 48*). We suggested that *Sulfurovum* as one of major primary producers in this sulfur-driven microbial ecosystem in IH stage of meromictic lake (Fig. 7).

### Complete holomictic stage

During CH in 2016, deep oxygenation expanded the oxic reservoir and depressed the redoxcline (>20 m) (*11*), resetting electron-donor/acceptor availability throughout the water column and coinciding with a decline in α-diversity. Co-occurrence networks simultaneously exhibited fewer positive edges and strengthened negative associations, consistent with enhanced niche partitioning and competitive exclusion as resources were re-allocated during collapse of vertical stratification. The bacterial community was dominated by *Campylobacteria*, *Gammaproteobacteria*, *Alphaproteobacteria*, and *Cyanobacteria*, with minor but persistent populations of sulfur related groups - CSB and PSB (Fig. 2D, Fig. 7). The coexistence of these lineages across the entire water column suggests that, even under whole-lake oxygenation, fine-scale micro-oxic and sulfidic niches likely persisted. Detection genes in metagenome which related to sulfur redox processes (*sqr*, *tsdA*, *soxABCDXYZ*, *soeABC*) and assimilatory nitrogen genes (*nasA/B*, *nirA*, *glnA*, *gdhA*, *ureC*) remained detectable at moderate normalized coverage (0.5–1.5 CPM; Fig. 6). It indicated a sustained potential for thiosulfate/sulfite oxidation and nitrogen uptake and recycling rather than denitrification. These genes encode the nitrate/nitrite assimilation and ammonium incorporation pathways, reflecting a shift toward nitrogen retention and regeneration under fully oxic yet nitrogen-limited conditions (*49*). Taxonomic composition of CPM contributions within sulfur redox genes (*sqr*, *tsdA*, *soxABCDXYZ*, *soeABC*) and assimilatory nitrogen genes (*nasA/B*, *nirA*, *glnA*, *gdhA*, *ureC*) were contributed predominantly by *Alphaproteobacteria* (notably *Rhodobacteraceae*-affiliated lineages such as *Yoonia* and related taxa) and *Gammaproteobacteria*, with additional contributions from Campylobacteria and Cyanobacteria (Suppl. Fig. S15 and Suppl. Fig. S16). This contributor profile broadly agrees with our 16S rRNA amplicon results, in which *Campylobacteria*, *Gammaproteobacteria*, *Alphaproteobacteria*, and *Cyanobacteria*.

Among six only sulfur-related MAGs in CH, *Yoonia* MAG (phylum *Pseudomonadota;* class *Alphaproteobacteria; Rhodobacterales; Rhodobacteraceae*) and UBA2463 MAG (phylum *Pseudomonadota*; class *Gammaproteobacteria*; order *Burkholderiales*; family *Burkholderiaceae*_A) were core sulfur-related MAGs. Core MAGs represent all stages retained capacities in supplied a stable metabolic backbone spanning all four stages (IH → CH → DM → M), with their relative contributions rebalanced during CH. Notably, *Yoonia* MAG encoded genes for assimilatory nitrate/nitrite reduction, whereas UBA2463 MAGs encoded nitrogen fixation genes (*50–52*). In our study, UBA2463 MAGs were rare (≤0.03%) and detected only in two oxic depths (DM_AOX, 23 m; M_OX, 2 m). In contrast, *Yoonia* maintained high relative abundance in total community (2-8%), especially increasing to 5% in CH stage. *Yoonia* is a genus within the *Rhodobacteraceae* that was initially isolated mainly from marine habitats (including algal-associated microhabitats and sediments) but has also been reported from brackish and lacustrine ecosystems (e.g., strains from Qinghai–Tibetan Plateau lake) (*50, 52, 53*). From previous studies, only a *Yoonia* sp. from plastic biofilms in the Great Pacific Garbage Patch has been reported to harbor a photosynthesis gene cluster (PGC) and to produce bacteriochlorophyll a (BChl *a*), indicating that some members may have aerobic anoxygenic phototrophic (AAP) potential (*54*). However, most species descriptions and genome reports do not document PGC or BChl genes, supporting the view that BChl is not a genus-wide conserved trait but rather a strain-or species-level variable. In our study, the *Yoonia* MAGs not only encompass the core metabolic repertoire described in the literature (extensive Sox system and nitrate/nitrite reduction) but also carry genes associated with bacteriochlorophyll, suggesting a possible AAP-derived phototrophic subsidy at oxic–microoxic interfaces. It indicated that this generalist aerobic bacteria are ubiquitous in the meromictic lake and especially dominate while the lake in holomictic condition. They, could replace anaerobic bacteria, constantly supported the fundamental nutrient cycle for meromictic lake in CH stage.

In addition, the genome information from CH-unique MAGs implied that they provided the short-term nutrient transformation and polymer-to-monomer carbon degradation functions. CH-unique MAGs highlight the processes that become especially important immediately after mixing Polymer-to-monomer carbon funnel. Among six CH-unique MAGs, three MAGs *Verrucomicrobiota* (*Chthoniobacterales*), *Planctomycetota* (*Pirellulales*), and *Bacteroidota* (*Kapabacteriales*) typically enriched in carbohydrate-active enzymes (CAZymes) (*55–57*). Especially, planctomycetes had sulfatases for degrading algal/gel-like sulfated polysaccharides (*58*). Thus, the apparent of these CH unique-MAGs might be related to the sharp increasing in phytoplankton in CH (*11*). It was consisted with previous paper that genes enrichment in CAZymes/sulfatases suggest oxygen-driven remineralization of algal-derived polysaccharides following the spring bloom (*59, 60*).

### Developing meromictic stage

As stratification re-established after CH, together with seasonal forcing, caused the redoxcline to shallow and sharpen, partitioning oxic and reduced waters once again. The microbial community structure followed this rebuilding of chemical niches. Many bacterial groups detected during CH persisted into DM, but the relative abundance shifted markedly, especially in DM_AOX.

After mixing, many residual small molecules likely derive from particle resuspension and upward transport, stress-induced cell lysis/exudation, extracellular depolymerization, and DOM, potential leaving methanol/formaldehyde–formate, methylamines, acetate and NH₄⁺ available as the water column re-stratifies based on metagenomic genes in bacterial metagenome. In DM_OX, higher gene abundance of formaldehyde→formate oxidation (*gfa*, *frmA/B* → *fdhA*, *fdoG/B/H/I*), methylamine (*mauA/B*, *mgs*/*mgd*), and nitrification (*hao*, *nxrA/B*) pathways, suggesting continued oxidation of residual small substrates and reduced nitrogen under aerobic condition. DM_OX also had many genes involved in carotenoid genes (*crt*) and the bacteriochlorophyll/light-harvesting suites (*bch*, *puh*, *puf*) and remained in M_OX. This indicated that the DM_OX probably induced the activity of aerobic anoxygenic phototrophs, which needed light harvesting to assist their heterotrophic processing in oxic layers (*61, 62*). Taxonomic composition of CPM contribution profiling indicates that this pattern was primarily driven by proteobacterial lineages consistent with AAP, notably *Yoonia*, UBA2463 sp018402635 (family *Burkholderiaceae*), *Oceanicaulis* and with minor contributions from *Rhodobacteraceae*-affiliated taxa (e.g., *Roseicyclus*, *Qipengyuania*) (Suppl. Fig. S18). In parallel, cyanobacterial taxa (e.g., *Cyanobium* and related lineages) contributed substantially to pigment-related transcripts, consistent with the concurrent presence of oxygenic phototrophs in oxic layers. Complementarily, 16S rRNA profiles showed an increase of purple sulfur bacteria (*Thiocapsa*) in the DO_OX and chemocline, enhance the contemporaneous reinforcement of the sulfur redox loop across the interface.

When rebuild of the stratification, the detection genes of unique MAGs in the anoxic layer suggested the return of sulfur-driven metabolism and low-oxygen nitrogen respiration. DM_AOX MAGs showed coherent sulfur modules—notably SQR (*sqr*) and flavocytochrome c sulfide dehydrogenase (*fccA/B*) for H₂S oxidation, Sox subunits (*soxA*/*B*/*X*/*Y*/*Z*/*C*/*G*) for thiosulfate oxidation, TsdA (*tsdA*) for thiosulfate → tetrathionate interconversion, and sulfide dehydorgenase (*soeA*/*B*/*C*) for sulfite oxidation, together with high gene detection in sulfur-relay/electron-bifurcation proteins (*Tus*/*DsrEFH*/*qmo*) of metagenome data in this study (*63, 64*). This set of genes in DM_AOX suggested a re-established chemolithoautotrophic scaffold at the redoxcline, potentially capable of processing reduced sulfur that diffuses upward from below or is regenerated on particles (*65*). At the same time, the enrichment of *hao* and *nxrA/B* genes points to the onset of micro-oxic nitrification, most likely occurring within particle-associated or near-interface niches where O₂ persists. The NO₂⁻ and NO₃⁻ generated *in situ* are probably consumed by co-occurring denitrification (*nirK/S*, *norB/C,nosZ*) and DNRA (*nrfAH*/*hcp*) pathway, forming a tightly coupled short-circuit of the nitrogen cycle (*66, 67*). While operation with the oxidizing reduced sulfur (*sqr*/*fcc*/*sox*/*tsdA*/*soe*), the electrons released might be used to nitrogen compounds (*68, 69*). Those electrons either reduce NO/N₂O to N₂ (denitrification) or turn NOx back into NH₄⁺ (DNRA). Because these ‘give-and-take’ reactions co-occur in the same micro-niches, it might be resulted taxa co-occur more and form tighter modules in DM_AOX.

### Meromictic stage

After a year for developing stable meromictic lake condition, meromixis was gradually established, the lake settles into a durable vertical arrangement in which the chemocline formed a steady redox environment. The oxic M_OX exhibited a pronounced genetic potential for sulfur oxidation, as reflected by the consistent detection of sulfur-oxidation genes across metagenomic assemblies. These patterns potentially indicate the presence of an oxically active sulfur-oxidizing community, most likely utilizing thiosulfate (S_2_O_3_^2-^) and other partially oxidized sulfur intermediates produced at the chemocline or on particles, with oxidation proceeding in the oxygenic surface (*70*). Notably, the chemocline zone also harbored abundant populations of purple sulfur bacteria (PSB, e.g., *Thiocapsa*), as supported by 16S rRNA profiles, indicating that anoxygenic phototrophs may contribute to sulfide and intermediate sulfur oxidation under low oxygen and light-exposed conditions. This sulfur-mediated phototrophy likely coexisted with photoheterotrophic and photoautotrophic phototrophy, together representing two modes. First, a direct phototrophic-sulfur oxidation mode where the groups harboring *soxA*/X/Y/Z and soeA/B/C are capable of oxidizing thiosulfate and sulfite while harvesting light (APP). From CPM contributions taxonomy profiling, they were *Rhodobacteraceae* (*Roseicyclus* and *Yoonia–Loktanella*) which had the aerobic anoxygenic phototroph signature (bacteriochlorophyll genes), as well as UBA2463 sp018402635, *Oceanicaulis*, and *Hyphomonas* (*71*). Second, an indirect coupling mode is suggested, in which cyanobacterial oxygenic photochemistry and particle-associated microgradients, it continuously generated to partially oxidized sulfur intermediates (e.g., S₂O₃²⁻, SO₃²⁻) that are utilized by co-occurring sulfur oxidizers in the low oxic place (*72*).

The anoxic M_AOX exhibited an enrichment of metagenomic *nif* genes, suggesting active N_2_ fixation under oxygen-depleted, sulfidic conditions where could combine nitrogen is scarce. Such anaerobic nitrogen fixation has been increasingly recognized in lake environments. In the same layer, H₂S concentrations gradually rose, aligning with *in situ* sulfate/sulfur reduction. At the same time, many M_AOX MAGs encode sulfur-redox electron-transfer mechanisms. It also indicated efficient routing of electrons from strongly reducing electron donors (H₂/formate) into a coupled reduction (H₂S production)–oxidation (intermediate consumption) loop (*73*). Consistently, the maximum relative abundance of SRB in 16S rRNA analysis can reached to 21% in M_AOX (October 2018). Four M_AOX unique-MAGs were sulfur-related bacteria: two of them were *Desulfobacteria* class, one of them was *Desulfobulbia*, and one of them was *Dethiobacteria*. They were obligately anaerobic, often halo/alkaliphile lineages frequently recovered from soda/saline lakes and sulfidic sediments (*74–76*). This guild likely supplies *in situ* sulfide at the chemocline and consumes fermentative H₂/formate, helping to close the reduction ↔ oxidation loop of the sulfur cycle within M_AOX micro-niches. Also, several bacterial MAGs harbored *cdh*, *mtaA/B/C*, and *mttB* genes; we interpret it consistent with the operation of the Wood–Ljungdahl pathway for bacterial acetogenesis and bacterial methylotrophy rather than archaeal methanogenesis (*77–80*). From above, the M_AOX co-occurrence network is comparatively sparse, showing fewer and weaker edges than other stages—consistent with strong environmental filtering and tightly specialized micro-niches that limit broad co-variation among taxa.

## Conclusion

Complete overturns are rare in land-locked meromictic systems, where strong density gradients usually prevent full circulation. Through fully dissected the relative abundance and metagenome in bacterial community in every depth and hydrodynamic stage of Lake Shira, microbial assemblages reorganized across intermittent holomixis, complete holomixis, developing meromixis, and stable meromictic lake stage. Each stage reflects a distinct restructuring of energy fluxes, redox processes, and functional capacities, collectively shaping the composition of microbial succession in meromictic Lake Shira.

During the short transition IH stage, syntrophy-driven associations were enhanced in the anoxic layer, highlighting the activity of anoxic bacteria such as sulfur-oxidizing bacteria (*Sulfurovum*) as early fundamental producers. In contrast, the CH stage reset the entire water column, leading to homogenization of niches but functional reweighting rather than extensive taxonomic replacement. Core MAGs might maintain sulfur and nitrogen pathways, while CH-unique taxa temporarily fueled carbon turnover within micro-oxic interfaces. As stratification re-established in DM, niche partitioning was enriched. DM_OX exhibited elevated capacities for oxidation of residual C1 substrates and reduced nitrogen, and pigment/light-harvesting genes point to increased photoheterotrophic/AAP activity, driven by a mixture of cyanobacterial phototrophs and proteobacterial AAP candidates. Finally, the stable M stage formed a dual system—surface oxic layers supporting photoautotrophic and photoheterotrophic modes, and anoxic zones dominated by sulfate reducers and diazotrophs, together closing the sulfur and nitrogen loops through tightly specialized groups.

## Acknowledgement

This work was supported by the National Science and Technology Council (NSTC; formerly the Ministry of Science and Technology, MOST), Taiwan (NSTC 113-2611-M-031-001 and NSTC 114-2611-M-031-001 to Ya-Fan Chan; MOST 110-2923-B-001-004-MY3 and MOST 106-2923-B-001-003-MY3 to Sen-Lin Tang), and by the State Assignment of the Ministry of Science and Higher Education of the Russian Federation (project No. FWES-2024-0024).

